# Mapping reward processing circuits: the orbitofrontal-nucleus accumbens white matter pathways and their associations to psychopathological traits

**DOI:** 10.64898/2026.10.05.756857

**Authors:** Santiago Mezzano, Fedal Saini, Veronica Orbecchi, Tobias Fernández-Borkel, Manu Raghavan, Mustafa Irfanoglu, Ahmad Beyh, Flavio Dell’Acqua

## Abstract

Dysfunctions in reward circuits, including projections from the orbitofrontal cortex (OFC) to the nucleus accumbens (NAcc), have been implicated in psychopathology. The OFC is functionally divided into the medial OFC (mOFC), which processes rewards, and the lateral OFC (lOFC), which responds to non-reward or punishment stimuli. We investigated the structural connectivity of the lOFC-NAcc and mOFC-NAcc pathways and their associations with psychopathological traits using tractography-defined ROI analysis. Diffusion-weighted imaging data from 165 participants (aged 22 - 35, 53% male) in the Human Connectome Project (HCP) were analysed using spherical deconvolution tractography. MegaTrack, a novel tool, enabled tract segmentation via a single manual dissection mapped back to native space. Virtual dissection identified four bilateral pathways connecting the lOFC and mOFC to the NAcc, from which fractional anisotropy (FA) was extracted, and an lOFC/mOFC ratio was computed. Psychopathological traits were measured using HCP Achenbach Adult Self-Report scores. The lOFC-NAcc pathway showed positive associations with psychopathology and the mOFC-NAcc pathway negative associations. The lOFC/mOFC ratio emerged as a stronger predictor than individual pathways, with significant associations for depression, somatic problems, anti-social personality traits, and inattention. Effects were lateralized to the left hemisphere. These findings show that the lOFC-NAcc and mOFC-NAcc pathways can be structurally separated with tractography and display opposing correlations with psychopathological traits in a healthy population. Their ratio may index reward-processing dysfunction, and future research should test whether it is sensitive to dysfunction in clinical populations.

## Introduction

Reward-circuitry dysfunctions involving the orbitofrontal cortex (OFC) have been implicated in the pathophysiology of numerous debilitating conditions, including major depressive disorder (Cui, 2024), anxiety disorders (Milad et al., 2007), substance use (Bracht et al., 2021), obsessive-compulsive disorder (Abe, 2015), and eating disorders (Frank, 2013). Recent findings emphasise a functional and structural distinction between the lateral (lOFC) and medial (mOFC) divisions of the OFC (Rolls, 2019; Hsu, 2020). The lOFC is involved in processing unpleasant stimuli (punishments) and unfulfilled reward expectations (non-rewards) (Rolls et al., 1994; Nogueira et al., 2017; Rolls, 2020), whereas the mOFC is activated by pleasant stimuli and rewards (Grabenhorst et al., 2011; Rolls, 2019; Xie et al., 2021). These differences are supported by converging evidence from neuroimaging (Rolls, 2020), behavioural (Wang, 2023), post-mortem (Henssen, 2016), chemogenetic (Oyama, 2022), and optogenetic (Winkowski, 2017) studies. Most recently, intracranial recordings in humans revealed opposing activity in the two divisions during approach–avoidance decisions, with mOFC activity increasing and lOFC activity decreasing prior to choosing to approach, alongside anticorrelated dynamics consistent with competitive interactions between them (Starkweather et al., 2026). Accordingly, lOFC and mOFC dysfunctions appear to contribute to distinct alterations in reward processing and, in turn, to the emergence of psychopathological traits.

These distinctions have prompted numerous investigations into their roles across clinical populations using functional neuroimaging. For example, a review by Zhang et al. (2016) highlights differences in resting-state activation of the OFC between patients with major depressive disorder (MDD) and healthy controls, reporting under-activation of the mOFC and over-activation of the lOFC in MDD. Similarly, Xie et al. (2021) demonstrated reduced mOFC activation during reward-related tasks alongside increased lOFC activation during unmet reward expectations in MDD, effects suggested to be modulated by antidepressant medication (Rolls et al., 2019; Wang et al., 2023).

Further studies identify abnormal activation and functional connectivity of the nucleus accumbens (NAcc) across psychopathologies. The NAcc lies in the ventral striatum, where many of the brain’s reward systems converge (Sesack, 2010), and is a key node in cortico-basal ganglia loops, receiving inputs from cortical regions including the OFC (Mobini, 2002; FitzGerald, 2009). Neuronal recordings in primates and functional imaging suggest that the identity of stimuli afferent to the OFC is represented independently of their reward value (Rolls et al., 2005); that is, reward information does not influence how the OFC encodes the descriptive features of incoming stimuli. Instead, reward-value representations that guide behaviour are conveyed through efferent cortical projections to subcortical regions, including glutamatergic pathways to the NAcc (Rushworth et al., 2012; Rolls, 2019). The OFC thus provides a route for reward-related information to reach the NAcc, where it interacts with dopaminergic inputs from the ventral tegmental area (Lindenbach, 2022; Schultz, 2016 ), a process essential for decision-making and goal-directed behaviour (Haber, 2010).

White matter (WM) projections from the OFC to the NAcc are central to reward circuitry, and dysfunctions in these connections have been associated with psychopathology. Increased mOFC-NAcc WM connectivity has been linked to mania/hypomania (Damme et al., 2017), consistent with a reward-hypersensitivity hypothesis (Sazhin, 2024), while reductions in NAcc volume, FA, and tract length of the OFC-NAcc network have been linked to craving in alcohol use disorder (Bracht et al., 2021). However, no study has investigated and compared the mOFC-NAcc and lOFC-NAcc pathways independently.

The present study adopts a more targeted approach by clearly distinguishing the lOFC and mOFC (Rolls, 2014), which have been shown to differ in terms of cytoarchitecture (Henssen, 2016), function (Kahnt, 2012), and their patterns of WM projection to the striatum (Haber, 2017). Consistent with this, recent tractography analyses have reported direct differences in the fibre bundles of the mOFC-NAcc and lOFC-NAcc, suggesting a divergence in functional specialisation related to learning and decision-making (Hsu, 2020), and highlighting the need to consider these subregions independently.

According to the non-reward attractor theory of depression (Rolls, 2016), emotions are framed as states elicited by rewards, their omission (non-reward), or punishments (Rolls, 2023), with the lOFC and mOFC playing distinct roles in this process. Within this framework, the lOFC is proposed to sustain attractor states, in which persistent firing in recurrently connected populations encodes non-reward and punishment (Khona, 2022), with activity lasting several seconds after negative stimuli (Rolls, 2021). According to this model, such non-reward attractors are thought to become overactive in depression, producing the prolonged negative states that characterise negative emotions, while reward-related mOFC responses are correspondingly hindered.

Critically, the theory suggests that reward processing in the OFC operates as an opponent balance between a reward-sensitive mOFC and a punishment-sensitive lOFC, such that pathology reflects a shift in their relative weighting rather than a change in either region alone. Because these cortical signals reach the NAcc through segregated WM projections (Haber, 2016), and dopaminergic reinforcement can drive structural change in these tracts over time (Koch, 2013), such a functional imbalance may be mirrored in the relative microstructure of the two pathways. This shift in relative weighting could be captured by a single index: a ratio between lOFC-NAcc and mOFC-NAcc reflecting reward dysfunction across the punishment and reward limbs of the circuit.

This study builds on current research in three ways. First, it examines OFC-NAcc structural connectivity using diffusion-weighted imaging from the Human Connectome Project (HCP) S1200 release (Van Essen et al., 2013), applying spherical deconvolution tractography (Dell’Acqua and Tournier, 2019) to identify distinct WM projections linking the mOFC and lOFC to the NAcc. Second, it examines how these pathways relate to psychopathological traits measured with the Achenbach Adult Self-Report (Achenbach & Rescorla, 2003), providing insight into reward-network dysfunction and its association with mental health. Third, alongside individual tract metrics, it introduces a quantitative metric, the lOFC/mOFC ratio, to test whether imbalances between the OFC subdivisions and their NAcc projections are better characterised by the complementary interaction of the two tracts than by either tract alone.

We hypothesise that the excessive sensitivity to punishment and insufficient sensitivity to reward characterising depression and related psychopathologies may be expressed as WM microstructural differences, specifically greater FA in lOFC-NAcc pathways and lower FA in mOFC-NAcc pathways. As these mechanisms are inherently dimensional, we test whether this pattern is detectable across subclinical variation in psychopathological traits within a healthy population

## Materials and Methods

### Participants

High-resolution diffusion and structural data from 200 healthy, right-handed young adults (Oldfield, 1971) (100 male, 100 female; aged 22-35 years) were drawn from the Human Connectome Project (HCP) S1200 release (Van Essen et al., 2013; http://www.humanconnectome.org). Participants were screened to confirm the absence of familial relationships and of any history of neurological or psychiatric disorder. Data was used in accordance with the HCP Open Access and Restricted Data Use Terms.

### Image acquisition

Acquisition procedures have been described in detail elsewhere (Sotiropoulos et al., 2013; Van Essen et al., 2013) and are summarised briefly here. Data were acquired on a customised 3T “Connectome Skyra” scanner (Siemens Healthineers, Erlangen, Germany) at the Mallinckrodt Institute of Radiology, Washington University, St. Louis, MO, USA. Diffusion-weighted images were obtained with a multiband echo-planar imaging sequence (multiband factor 3; 1.25 mm isotropic voxels; field of view 210 × 180 mm; 111 slices; TE 89.5 ms; TR 5520 ms) across three shells (b = 1000, 2000, and 3000 s/mm^2^). Each session comprised six runs from three gradient tables, each table containing six non-diffusion-weighted (b = 0) volumes and 90 diffusion-weighted directions, acquired in both left-right and right-left phase-encoding directions for susceptibility-distortion correction.

### Image processing

The OFC is a challenging region for diffusion imaging because its proximity to air-filled sinuses produces strong field inhomogeneities, causing EPI distortion and signal dropout in inferior frontal regions (Devlin et al., 2000). Given the small calibre and high inter-individual variability of the lOFC/mOFC-NAcc pathways, reconstructions here are particularly susceptible to signal loss and premature streamline termination. To improve robustness, the HCP diffusion data were reprocessed with a customised pipeline in TORTOISE v3.2 (Pierpaoli et al., 2010).

First, data was denoised using the Marchenko-Pastur principal component analysis approach, which identifies and removes noise-only components by exploiting redundancy in the diffusion signal (Veraart et al., 2016). Gibbs-ringing artefacts were corrected via local sub-voxel shifts (Kellner et al., 2016). Motion and eddy-current distortions were corrected with the DIFF_PREP module (Rohde et al., 2004), and susceptibility-induced distortions with the DR-BUDDI framework (Irfanoglu et al., 2015), which estimates the displacement field from images acquired with opposing phase encoding and incorporates registration to the undistorted structural T2-weighted image. Additionally, gradient non-linearity and B0 drift were corrected using in-house software.

### Model fitting, tractography and MegaTrack segmentation

Spherical-deconvolution tractography was performed in StarTrack (http://nbl-research.github.io) using the damped Richardson-Lucy algorithm (Dell’Acqua et al., 2010), with fibre response α = 1.5, 200 iterations, and regularisation parameters η = 0.0015 and ν = 16. Streamlines were propagated with a deterministic Euler-like algorithm with a 45° angle threshold, an absolute fODF amplitude threshold of 0.003, and a 1 mm step size; only streamlines 20-300 mm in length were retained for dissection (Dell’Acqua et al., 2013).

Tracts were segmented using MegaTrack (Dell’Acqua, 2025), a novel efficient technique that merges tractography from a group of subjects into a common standard space so that a single dissection can be applied across the entire sample. Because all dissections are performed in standard space, this approach yields unbiased, consistent segmentation and simplifies identification of anatomical landmarks. Following the standard MegaTrack procedure, affine and non-linear transformations to MNI space were computed with ANTs (Avants et al., 2011) from anisotropic power maps (Dell’Acqua et al., 2014) and applied to all streamlines. Streamline compression was applied with a maximum error of 0.4 mm. The merged tractography dataset was dissected in TrackVis (Wang et al., 2007). Each streamline was tagged with both a subject and a unique streamline identifier, allowing surviving streamlines to be traced back to each subject’s native space for metric extraction.

### Virtual dissection, tract criteria and quality control

Virtual dissections used cortical and subcortical regions of interest (ROIs) for the medial OFC, lateral OFC, and nucleus accumbens (NAcc) derived from the FreeSurfer Desikan-Killiany parcellation (Desikan et al., 2006). Four pathways were reconstructed, connecting the NAcc to the lOFC and to the mOFC in each hemisphere (1 subcortical ROI × 2 cortical ROIs × 2 hemispheres). Cortical and subcortical ROIs were defined as the start and end points of each tract, retaining only streamlines that terminated in both grey-matter ROIs (“either end” criterion). Spurious streamlines were removed with tract-specific exclusion (“not”) ROIs that discarded inter-hemispheric crossings, looping streamlines, and streamlines deviating substantially from the main bundle.

From each reconstructed tract, fractional anisotropy (FA) was extracted as the microstructural metric of interest. FA is a rotationally invariant diffusion-tensor metric quantifying the directional coherence of water diffusion, enabling orientation-independent comparison across subjects (Basser et al., 1996); it is sensitive to along-tract variation in WM microstructure, including myelination, axonal integrity, and fibre dispersion.

Given the vulnerability of inferior frontal regions to signal dropout strict quality control was applied to maximise the validity of connectivity estimates: participants with fewer than four streamlines in any of the four tracts of interest were excluded. This conservative threshold minimises the influence of poorly reconstructed tracts. Together with one exclusion for missing behavioural data, this criterion removed 35 of the 200 participants, yielding a final sample of 165.

### Assessment of behavioural and emotional functioning

Participants completed the Achenbach Adult Self-Report (ASR), a 126-item questionnaire assessing behavioural, emotional, and adaptive functioning in adults aged 18-59 (Achenbach & Rescorla, 2003). The ASR provides a global index of functioning alongside scores for specific domains. The ASR total T-score (ASR_Top_T) was used as an aggregate index of overall behavioural and emotional functioning. In addition, six domain scores derived from the total ASR questionnaire were examined : Depression, Anxiety, Somatic Problems, Inattention, Hyperactivity, and Antisocial Traits. Although the HCP cohort comprises healthy adults without a history of psychiatric or neurological disorder, this multidimensional instrument captures inter-individual variation in subclinical behavioural and emotional traits across domains.

### Statistical analysis

Analyses were conducted in Python 3.11. Distributional normality was assessed and confirmed with Shapiro-Wilk tests. We first examined associations between individual tract FA values and total psychopathology (ASR_Top_T). To assess the stability of these associations, a repeated split-half validation was performed: the sample was randomly partitioned into two equal halves 30 times, partial correlations were computed independently within each half, and the proportion of splits in which both halves showed the same directional effect was recorded as an index of consistency.

Motivated by the opposing functional roles of the lateral and medial OFC-NAcc pathways, an lOFC/mOFC FA ratio was computed for each participant by dividing lOFC-NAcc FA by mOFC-NAcc FA, providing a single, self-normalised index of the relative balance between the two pathways. After examining the association between this ratio and total psychopathology (ASR_Top_T), subsidiary analyses tested associations between the ratio and each of the six ASR domain subscales. Correlations of subscales with individual lOFC and mOFC tracts are reported to compare against ratio and evaluate which tract is contributing more to the effect \. All correlations covaried for age and sex, and multiple comparisons were controlled using the Benjamini-Hochberg false discovery rate (FDR) procedure applied within each family of tests (Benjamini & Hochberg, 1995).

## Results

Our study analysed tract metrics from 165 participants acquired from the HCP (Van Essen et al, 2013). The sample’s mean age was 28.4 years (SD = 3.7, range 22-35), with a gender distribution of 47% female (n = 88). All participants are healthy young adults (aged 22-35) with no history of significant neurological or psychiatric disorders.

Preliminary analysis revealed a significant correlation between the age of participants and the average psychopathology score derived from the ASR (R = -0.21, p = 0.006), highlighting the need for controlling for age as a covariate in subsequent analyses (Supplementary Table 1). Additionally, we examined correlations among ASR subcategories to assess the shared variance between them (detailed in Supplementary Table 3), as these dimensional traits tend to co-occur and overlap in their item content, even at subclinical levels.

### Individual OFC-NAcc tract correlations with total psychopathology

Across the full sample, partial correlations (controlling for age and sex) between individual tract FA and total psychopathology (ASR_Top_T) revealed a consistent hemispheric pattern (Table 1). In the left hemisphere, the mOFC-NAcc tract was significantly negatively associated with total psychopathology (r = −0.177, p = 0.024), whereas the lOFC-NAcc tract showed a positive but non-significant association (r = 0.100, p = 0.204). Right-hemisphere tracts followed the same directional signs but were markedly weaker: the mOFC-NAcc tract showed only a weak negative trend (r = −0.095, p = 0.228) and the lOFC-NAcc tract essentially no association (r = 0.010, p = 0.898).

**Table 1.** Partial correlations between individual OFC-NAcc tract FA and total psychopathology (ASR total), controlling for age and sex. *p < 0. 05. Directional consistency is the proportion of 30 random split-half iterations in which both subsamples showed the same sign of correlation.

| Hemisphere | Tract | n | r | p | Directional consistency<br>(across 30 splits) |
| --- | --- | --- | --- | --- | --- |
| Left | mOFC-NAcc | 165 | -0.177 | 0.024* | 100% |
| Left | IOFC-NAcc | 165 | 0.100 | 0.204 | 83% |
| Right | mOFC-NAcc | 165 | -0.095 | 0.228 | 73% |
| Right | IOFC-NAcc | 165 | 0.010 | 0.898 | 7% |

Moreover, the two left-hemisphere tracts showed opposing and highly stable directions: the mOFC-NAcc association was negative in 100% of splits, while the lOFC-NAcc association was positive in 83%. Notably, although the left lOFC-NAcc correlation did not reach significance in the full sample, its positive direction - higher lateral FA with greater psychopathology - was reproduced in the large majority of subsamples, indicating a consistent if modest effect. Right-hemisphere directions were far less stable (mOFC-NAcc negative in 73% of splits, lOFC-NAcc positive in only 7%).

In summary, the opposing pattern, higher lateral OFC-NAcc FA associated with greater psychopathology, and higher medial OFC-NAcc FA with less, was a stable feature of the left-hemisphere tracts but unstable or absent on the right.

### The lOFC/mOFC FA ratio and total psychopathology

To capture the relative balance between the two pathways, we computed the l OFC/mOFC FA ratio. The left-hemisphere ratio was positively correlated with total psychopathology (r = 0.282, p < 0.001, p_FDR < 0.001), indicating that individuals with relatively greater lateral than medial OFC-NAcc FA exhibited higher overall psychopathological traits. This association was stronger than that of either constituent tract alone (Figure 1). The right-hemisphere ratio did not reach significance after FDR correction (r = 0.109, p_FDR = 0.162), mirroring the weaker and less stable right-hemisphere tract associations. Given this left-lateralisation, all post hoc-subsequent analyses focus on the left-hemisphere ratio.

**Figure 1.**
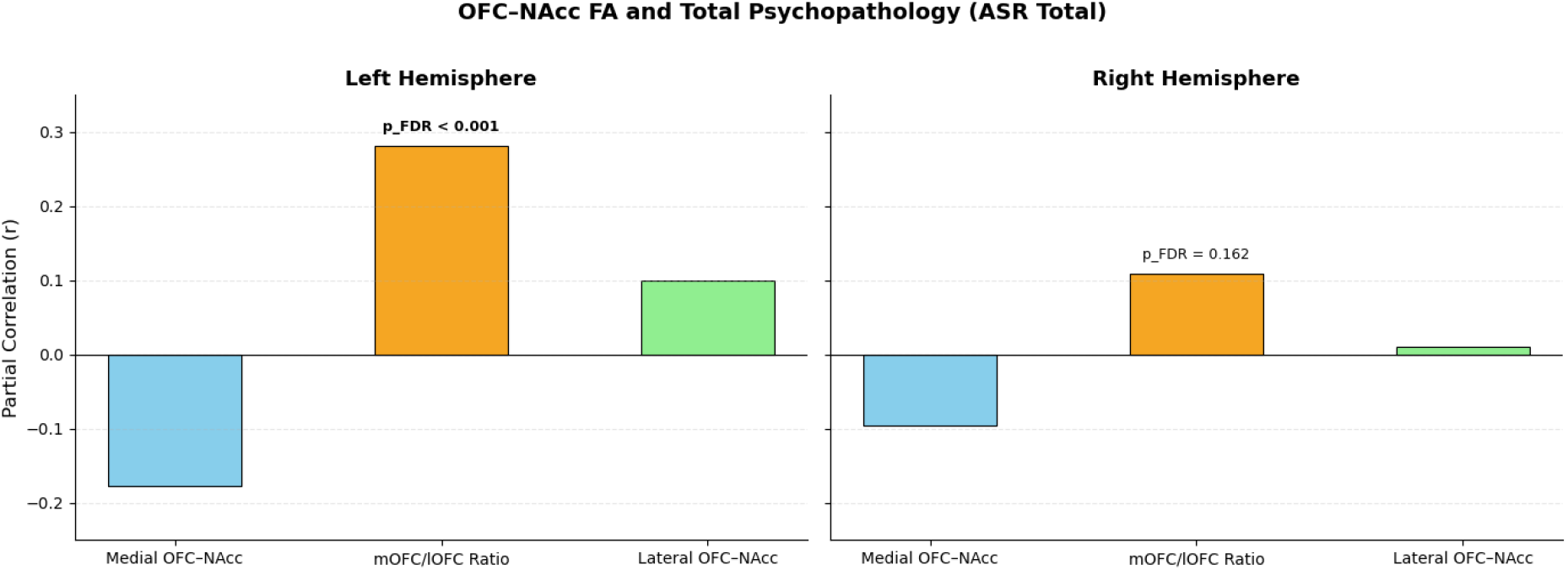
Partial correlations (controlling for age and sex) between OFC-NAcc FA and total psychopathology (ASR total) for the medial tract, the lateral tract, and the lOFC/mOFC ratio, shown separately by hemisphere. p_FDR annotated for the ratio.

### lOFC/mOFC FA ratio and specific psychopathological traits

We next examined how the left lOFC/mOFC ratio related to the six ASR domains (Figure 2; Supplementary Table S2). Four of the six domains were positively and significantly associated with the ratio after FDR correction: inattention (r = 0.273, p_FDR = 0.003), antisocial traits (r = 0.231, p_FDR = 0.007), depression (r = 0.229, p_FDR = 0.007), and somatic problems (r = 0.205, p_FDR = 0.013). Anxiety showed a trend-level association that did not survive correction (r = 0.159, p_FDR = 0.051), and hyperactivity showed no association (r = 0.049, p_FDR = 0.532). Across every domain, the ratio showed stronger associations than either the medial or lateral tract alone.

**Figure 2.**
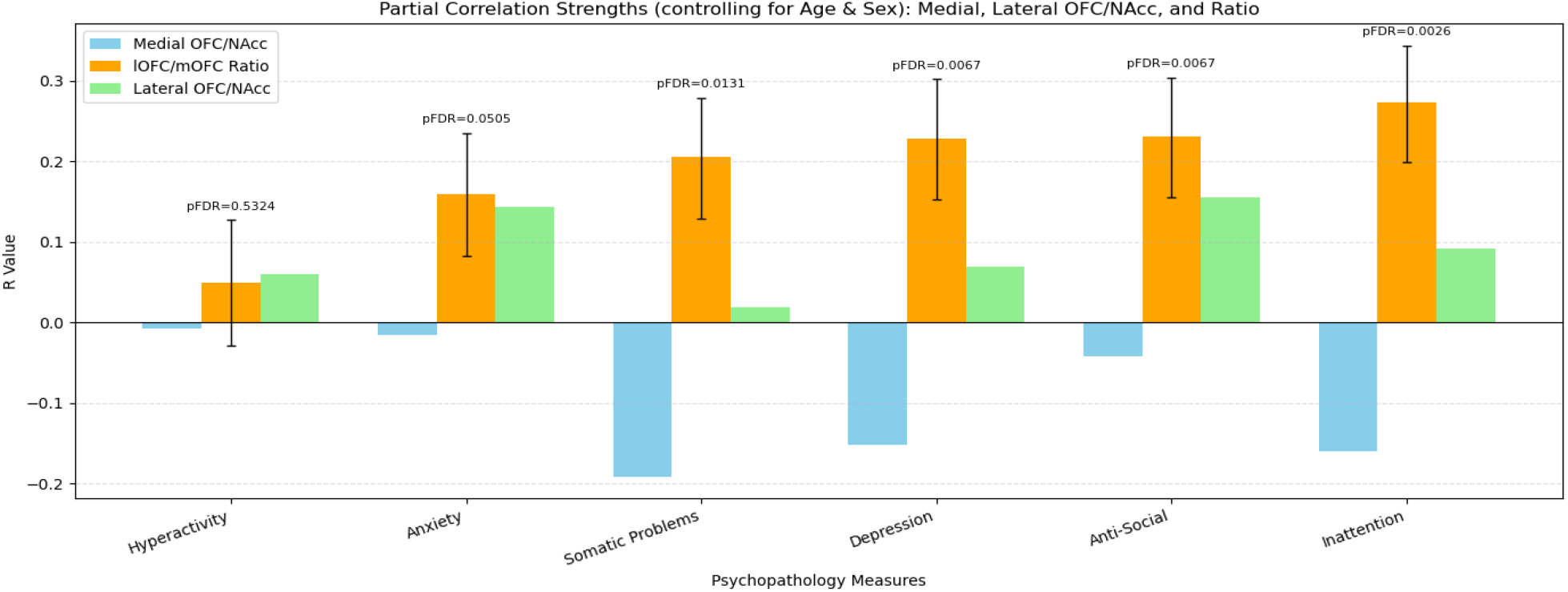
Partial correlations (controlling for age and sex) between the left lOFC/mOFC FA ratio and each ASR domain, with the medial and lateral tract correlations shown for comparison. Error bars are Fisher z-derived ±1 SE; p_FDR annotated per domain.

## Discussion

The present study examined whether OFC-NAcc WM microstructural organisation, indexed by tractography-derived fractional anisotropy (FA), is associated with psychopathological traits related to reward-system function in a healthy HCP sample. We focused on the distinction between lateral and medial OFC projections to the NAcc, testing whether inter-individual differences in lOFC-NAcc FA, mOFC-NAcc FA, and the ratio between them were associated with total psychopathology and with individual ASR symptom domains. Drawing on functional accounts of medial-lateral OFC differentiation supported by previous literature, we predicted that traits marked by oversensitivity to non-reward processing would correspond to higher lOFC-NAcc and lower mOFC-NAcc FA, and hence an elevated lOFC/mOFC ratio.

Three main findings emerged. First, lateral OFC-NAcc FA was positively associated with psychopathological traits, whereas medial OFC-NAcc FA was negatively associated. Second, the lOFC/mOFC FA ratio tracked psychopathology more robustly than either tract alone - both for the total score and, at the symptom level, for inattention, antisocial traits, depression, and somatic problems after FDR correction; anxiety showed a trend that did not survive correction, and hyperactivity showed no association. Third, these associations were left-lateralised, with no significant effects in the right hemisphere.

### OFC-NAcc imbalance and reward/non-reward processing

As hypothesised, the healthy HCP sample revealed that psychopathological trait scores were positively associated with lOFC-NAcc FA and negatively associated with mOFC-NAcc FA (Figure 1).

These findings extend functional accounts of lateral and medial OFC differentiation by showing that structural variation in their projections to the NAcc is associated with dimensional psychopathological traits even within a healthy sample, potentially reflecting WM plasticity in response to trait-related variation (Metzler-Baddeley 2017). Specifically, the positive association for lOFC-NAcc FA and the negative association for mOFC-NAcc FA suggest that trait burden may relate to a structural imbalance between pathways implicated in non-reward/punishment and in reward processing. This is consistent with the non-reward attractor theory of depression and related models of reward-processing dysfunction, which characterise the lOFC as more involved in non-reward processing and the mOFC in reward or positive-outcome processing (Camara, 2009; Weinstein, 2023; Rolls, 2022).

This study is also the first to show that combining the two pathways into a single lOFC/mOFC FA ratio yields a more robust correlation with psychopathological traits than either tract individually. The value of this ratio as a candidate biomarker of reward processing-related psychopathology warrants further investigation and validation across clinical datasets and acquisition parameters. These findings also highlight the importance of segregating the lateral and medial OFC as functionally distinct regions, rather than conflating them into a single functional unit (Kringelbach, 2005).

### Left lOCF/mOFC FA ratio and psychopathology domains

Consistent with our hypothesis, the left lOFC/mOFC FA ratio was associated with depressive traits. However, the symptom-level analyses indicate that the relevance of this ratio may extend beyond depression alone. Significant associations were also observed with inattention, antisocial traits, and somatic problems, whereas anxiety showed only a trend-level association This broader pattern suggests that the ratio may relate to variation across several psychopathological domains within a healthy sample rather than to depressive traits alone.

The strongest symptom-level association was with inattention, whereas hyperactivity showed no meaningful association. This dissociation suggests that the ratio may relate more closely to attentional or executive aspects of ADHD-like traits than to motoric hyperactivity. One possible interpretation is that this imbalance primarily affects goal-directed attention, rather than motor regulation more broadly. This would be consistent with the broader role of frontostriatal circuits in executive control, particularly in contexts where behaviour depends on reward expectations, effort allocation, or sensitivity to negative outcomes. Because the sample comprised healthy participants without clinical ADHD diagnoses, this should be framed as an association with dimensional inattention traits rather than as evidence for ADHD subtypes or differential diagnosis.

The association with somatic problems may also be read within a reward/non-reward framework, though more tentatively. The ASR somatic-problems scale captures physical complaints that may partly overlap with pain, interoceptive dysregulation, and the affective evaluation of bodily discomfort. Frontal and frontostriatal circuits, including the OFC, contribute to pain valuation and its modulation: monetary reward has been shown to reduce pain perception via mOFC activation (Becker et al., 2017), intracranial recordings have linked increased fronto-striatal connectivity to chronic pain states (Shirvalkar, 2023), and more broadly, OFC circuits have been implicated in pain expectation and subjective pain evaluation (Cerqueira-Nunes, 2024). The positive association between somatic problems and the ratio may therefore reflect a relative bias toward aversive-over reward-related circuitry. This interpretation remains speculative, as we did not directly measure pain, interoceptive processing, or clinically diagnosed somatic symptom disorder.

Overall, the symptom-level associations pattern suggests that the left lOFC/mOFC FA ratio relates to several dimensions of psychopathological trait variation within a healthy sample. Depression provides the clearest link to the study’s reward/non-reward framework, while the stronger association with inattention suggests that OFC-NAcc pathway balance may also be relevant to attentional and executive-control traits. Future research should test whether these associations replicate in independent datasets and in clinically diagnosed populations, including depression, ADHD, somatic symptom disorders, and other conditions involving altered reward or punishment processing.

Although the ratio provided the most robust index of psychopathological trait variation, decomposing it into its constituent pathways suggested that the two tracts may contribute differentially across domains. Anxiety and antisocial traits were numerically driven by the lateral (lOFC–NAcc) pathway, whereas somatic problems, depressive traits, and inattention were more associated with the medial (mOFC–NAcc) pathway. Within the reward/non-reward framework, this grouping is coherent: anxiety and antisocial traits both involve altered sensitivity to threat, punishment, or aversive outcomes (Grupe & Nitschke, 2013; Gregory et al., 2015) functions attributed to the punishment-sensitive lOFC, whereas somatic, depressive, and inattentive traits may relate more to diminished reward or anhedonia (Pizzagalli et al., 2009; Borsini et al., 2020; Serretti, 2025: Ma, 2024), attributed more to the mOFC.

### Lateralisation of OFC-NAcc effects

The results also suggested a left-lateralised association between OFC-NAcc pathway balance and psychopathological traits: the left lOFC/mOFC FA ratio was significantly associated with total psychopathology, whereas the right-hemisphere ratio did not survive correction. This asymmetry is unlikely to reflect asymmetric tract reconstruction, as prior tractography mapping of the mOFC- and lOFC-NAcc pathways reported symmetric connectivity (Cartmell,2019). The relationship between relative lateral-medial OFC-NAcc microstructure and trait variation therefore appears to be more pronounced on the left hemisphere for reward processing.

Work on OFC lateralisation remains limited, particularly regarding reward processing and psychopathology. Some functional studies suggest that reward valuation and OFC connectivity show hemispheric asymmetries (López-Persem, 2020), but their anatomical and clinical significance is unclear. The present left-lateralised association should thus be treated as an exploratory observation requiring replication, rather than as evidence for established hemispheric specialisation in OFC-NAcc reward circuitry.

### Clinical relevance

These associations were obtained in a healthy sample and describe subclinical trait variation; any clinical implications are therefore indirect and require validation in patient populations. With that caveat, linking distinct OFC-NAcc pathways to reduced reward sensitivity and heightened punishment sensitivity has potential implications for behavioural, pharmacological, and neurostimulation approaches. The findings are consistent with the rationale underlying interventions such as Behavioural Activation Therapy (Karimpour-Vazifehkhorani, 2020), which promotes engagement with positive reinforcement and has been shown to particularly benefit individuals with reward processing-related symptoms including apathy (Jung, 2024); such approaches may, in principle, strengthen reward-related mOFC-NAcc circuitry. Conversely, therapies targeting sensitivity to aversive or non-rewarding stimuli, such as exposure therapy (Papalini, 2019) or dialectical behaviour therapy (Chapman, 2006), might be framed as dampening lOFC-NAcc involvement. Neurostimulation studies point in the same direction: transcranial magnetic stimulation disrupting lOFC activity has shown promise in depression (Feffer et al., 2018; Downar, 2019), while deep brain stimulation of the mOFC has been reported to improve mood (Rao, 2018). As tractography-guided Deep Brain Stimulation enables individualised targeting of specific pathways (Chan, 2024), our findings highlight the value of distinguishing lateral from medial OFC projections when planning such interventions. The lOFC/mOFC ratio may therefore offer a compact index for this purpose, though its clinical utility remains to be formally established.

### Limitations

Few limitations of the study need to be reported. First, FA is a non-specific microstructural index: although it may be sensitive to properties such as myelination, axonal density, and fibre coherence, it does not measure WM “integrity” directly, especially within voxels containing crossing or dispersing fibres. These associations should therefore be interpreted at the level of microstructural organisation, rather than as evidence for any specific biological mechanism. Second, due to its inferior frontal location, the OFC is especially vulnerable to susceptibility-induced distortion and signal dropout; despite dedicated preprocessing and strict streamline-count quality control, residual reconstruction errors cannot be completely excluded, and the weaker effects observed within the right-hemisphere may partly reflect these acquisition-related artifacts rather than biology. Moreover, the sample comprised healthy young adults, so the observed trait variation is subclinical and of restricted range. Lastly, the lOFC/mOFC ratio, while parsimonious, is a derived measure whose value depends on both constituent tracts; equal ratios can arise from different underlying configurations, and should therefore be reported alongside its individual constituent pathways rather than in place of them.

### Future directions

Future tractography studies should map the broader reward circuitry to further probe its relationship with psychopathological traits. The supracallosal anterior cingulate cortex, responsive to aversive stimuli (Grabenhorst & Rolls, 2011; Rolls, 2014), is structurally connected to the lateral OFC, whereas the pregenual cingulate cortex, activated by rewarding stimuli, is structurally and functionally connected to the medial OFC (Rolls et al., 2019; Hsu et al., 2020; Du et al., 2020). Extending analyses to additional pathways - such as basolateral amygdala-OFC connections (Lichtenberg, 2017) and mesolimbic projections - would help clarify the dynamics of the reward network and its relevance to psychopathology. Above all, testing whether the lOFC/mOFC ratio is sensitive to reward-system dysfunction in clinically diagnosed populations, and whether it carries predictive or treatment-relevant information, is the key next step.

## Conclusion

Taken together, these findings support the theoretical proposal that the lateral and medial OFC-NAcc pathways make distinct, opposing contributions to reward processing. By separating the two pathways with tractography, we show that their microstructure relates in opposite directions to psychopathological traits in a healthy population, with effects most pronounced in the left hemisphere. Crucially, the lOFC/mOFC FA ratio, which captures the balance between the two pathways, was more strongly associated with psychopathology than either pathway alone, and these associations extended beyond depressive symptoms to reward processing dysfunctions. The ratio therefore offers a structural index of the balance between non-reward/punishment and reward processing proposed by Rolls’ model of depression, and a means of testing this framework transdiagnostically, across conventional nosological boundaries. That these associations are detectable in individuals without clinical diagnoses suggests the measure is sensitive to subclinical variation, raising the possibility of identifying individuals at elevated risk before pathology emerges.

Replication in independent, longitudinal and clinical cohorts will be needed to establish whether the ratio can inform early identification and preventive or targeted interventions.

